# Mutations at the Alphavirus E2/E1 inter-dimer interface have host-specific phenotypes

**DOI:** 10.1101/2021.09.02.458808

**Authors:** Sophia C. Ren, Shefah A. Qazi, Brian Towell, Joseph CY Wang, Suchetana Mukhopadhyay

**Affiliations:** Department of Biology, Indiana University, Bloomington, IN 47405; Molecular and Cellular Biochemistry, Indiana University, Bloomington, IN 47405; Department of Microbiology and Immunology, Penn State College of Medicine, Hershey, PA 17033

## Abstract

Alphaviruses are enveloped viruses transmitted by arthropod vectors to vertebrate hosts. The surface of the virion contains 80 glycoprotein spikes embedded in the membrane and these spikes mediate attachment to the host cell and initiate viral fusion. Each spike consists of a trimer of E2-E1 heterodimers. These heterodimers interact at two interfaces: (1) the intra-dimer interactions between E2 and E1 of the same heterodimer, and (2) the inter-dimer interactions between E2 of one heterodimer and E1 of the adjacent heterodimer (E1’). We hypothesized that the inter-dimer interactions are essential for trimerization of the E2-E1 heterodimers into a functional spike. In this work, we made a mutant virus (CPB) where we replaced six inter-dimeric residues in the E2 protein of Sindbis virus (WT SINV) with those from the E2 protein from chikungunya virus, and studied its effect in both mammalian and mosquito cell lines. CPB produced fewer infectious particles in mammalian cells than in mosquito cells, relative to WT SINV. When CPB virus was purified from mammalian cells, particles showed reduced amounts of glycoproteins relative to capsid protein, and contained defects in particle morphology compared to virus derived from mosquito cells. Using cryo-EM, we determined that the spikes of CPB had a different conformation than WT SINV. Last, we identified two revertants, E2-H333N and E1-S247L, that restored particle growth and assembly to different degrees. We conclude the inter-dimer interface is critical for spike trimerization and is a novel target for potential antiviral drug design.

**IMPORTANCE:** Alphaviruses, which can cause disease when spread to humans by mosquitoes, have been classified as an emerging pathogen, with infections occurring worldwide. The spikes on the surface of the alphavirus particle are absolutely required for the virus to enter a new host cell and initiate an infection. Using a structure-guided approach, we made a mutant virus that alters spike assembly in mammalian cells but not mosquito cells. This is important because it identifies a region in the spike that could be a target for antiviral drug design.

## INTRODUCTION

Alphaviruses are most commonly transmitted by arthropod vectors, usually mosquitoes, to vertebrate hosts, including humans, birds, and horses (1, 2). While a majority of alphaviruses have an arthropod vector, a group of alphaviruses have been identified to transmit only between invertebrates (3), and others use a different vector to infect aquatic animals (4). Therefore, for an alphavirus to complete its infection cycle, it must be able to assemble particles in all of these host environments. Virus infection must rely on different host factors since the same viral proteins are synthesized.

The alphavirus genome consists of four nonstructural proteins and six structural proteins, which are required for viral genome replication and particle assembly, respectively (1, 2, 5, 6). The alphavirus particle consists of an inner nucleocapsid core, a host-derived lipid membrane, and 80 trimeric spikes on the surface of the virion (7). The spikes are trimers of E2-E1 heterodimers with each heterodimer forming the edge of a triangular spike. Both the E2 and E1 proteins each contain a single transmembrane domain and are embedded within the viral membrane. The endodomain of E2 interacts with the capsid protein in the core in a 1:1 ratio (8-12). Thus, the E2 protein transits the entire particle and helps align the core and the spikes, a unique feature of alphaviruses compared to other enveloped viruses (7).

Alphavirus spike assembly is a highly regulated process that depends on specific interactions between the viral proteins and host factors. The structural proteins are translated as a capsid-E3-E2-6K-E1 polyprotein (or capsid-E3-E2-TF when frameshifting occurs). Capsid is autoproteolytically cleaved from the rest of the polyprotein (13). E3 serves as the signal sequence to translocate the rest of the polyprotein into the endoplasmic reticulum (ER) (14, 15), where the cellular enzyme signalase cleaves the polyprotein into E3-E2 (also known as pE2 or P62), 6K, and E1 (16, 17). When programmed ribosomal frameshifting occurs the protein pE2 and TF are translated (5). In mammalian cells, the ER chaperones Erp57 and calnexin/calreticulin regulate the folding and disulfide bond formation of E2 (18-20), and BiP and protein disulfide isomerase do the same for E1 (18-22). pE2 and E1 form heterodimers within the ER, before transiting to the Golgi where trimerization of the stable spike complex is predicted to occur. Glycosylation of E2 and E1 occur in the ER and Golgi. In the ER and Golgi, there is a slight decrease in pH and the E3 proteins acts as a clamp and prevents low pH mediated dissociation of the E2-E1 heterodimer (23-25). These spikes in the stable conformation are transported to the plasma membrane through the host secretory system. In the late secretory pathway, the cellular enzyme furin cleaves E3 from E2, which acts as a priming event and converts the spike from the stable to metastable conformation (26, 27). At the plasma membrane, E2 interacts with the nucleocapsid core, and initiates budding and virus release. Particles may also bud from glycoprotein-containing vesicles called cytopathic vesicles II, and this pathway is used more by mosquito cells (28).

Voss et al. solved the Chikungunya virus (CHIKV) E2/E1 heterodimer in the metastable conformation and in complex with E3 at neutral pH in the stable conformation (29), and Li et al. solved the Sindbis virus (SINV) E2/E1 heterodimer at acidic pH (30). These structures showed the intra-dimer contacts within a heterodimer. Both groups fit the atomic heterodimers into cryo-EM structures of alphavirus virions to identify inter-dimer interactions, or contacts between heterodimers within the spike, and between the E2 and nucleocapsid core. E1 consists of three domains: I-III. Domain II contains the fusion peptide and makes extended contacts with E2 in the intra-dimer heterodimer (29-31). This intra-dimer interface contains the acid-sensitive region and has been studied in regard to fusion regulation and mutations that expand vector range (32, 33). E2 has three domains: A-C. Domain B is the most distal domain and acts as a cap of the distal end of E1 protecting the fusion peptide (29, 30). In the low pH structure by Li et al., Domain B was disordered suggesting it is the first portion of the heterodimer to undergo conformational changes in response to low pH (30). Domain A in E2 is the central domain and Domain C is the closest to the lipid bilayer. Domain C is sandwiched between Domain II of E1 from its intra-dimer heterodimer and the Domain II of E1 from the adjacent heterodimer, or E1’. Both Voss et al. and Li et al. identified residues in E2 Domain C that contact residues in Domain II of E1’ (29, 30). Based on these two structures, we hypothesized that E2 Domain C plays a key role in spike assembly, and disrupting the inter-dimer contacts between E2 and E1’ could affect trimer formation.

To test the role of Domain C in assembly, we substituted six amino acids in SINV E2 Domain C to the corresponding residues in CHIKV, an alphavirus in a different clade (2, 34). The E2 residues that were mutated are predicted to interact with E1’ residues in the adjacent dimer, or to “piggyback” on the dimers within a spike, and hence our mutant was named Chikungunya Piggyback (CPB). We determined CPB had slower growth and smaller plaques when grown in mammalian cells but not mosquito cells. CPB quickly gained second-site revertants, E2-H333N and E1-S247L, which were each able to restore growth in CPB. CPB grown in mammalian cells form particles with assembly defects but the revertants restored their assembly to various degrees. Further analysis of the spike conformations on the CPB by cryo-EM showed the spikes in CPB were more flexible than the spikes in WT virus.

## RESULTS

### Domain C of E2 is important for spike trimerization

The atomic structure of the CHIKV E2/E1 heterodimer identified the interface between E2 and E1 within a heterodimer and the intra-dimer contacts that were present. When these heterodimers were placed into the cryo-EM density of intact SINV virions, the potential contacts between heterodimers, or inter-dimer contacts, were identified (Figure 1A-1C). In the virion, Domain C of the E2 protein was sandwiched between two E1 proteins, one in the same heterodimer, designated as E1, and one from the adjacent heterodimer, designated as E1’ (Figure 1D, 1E) (29). We have colored the different domains of E1 and E2 in Figure 1E to illustrate these interactions of Domain C of E2. We hypothesized that Domain C would be important for spike trimerization because these inter-dimer contacts would bridge or connect the individual E2-E1 dimers into trimers.

**Figure 1:**
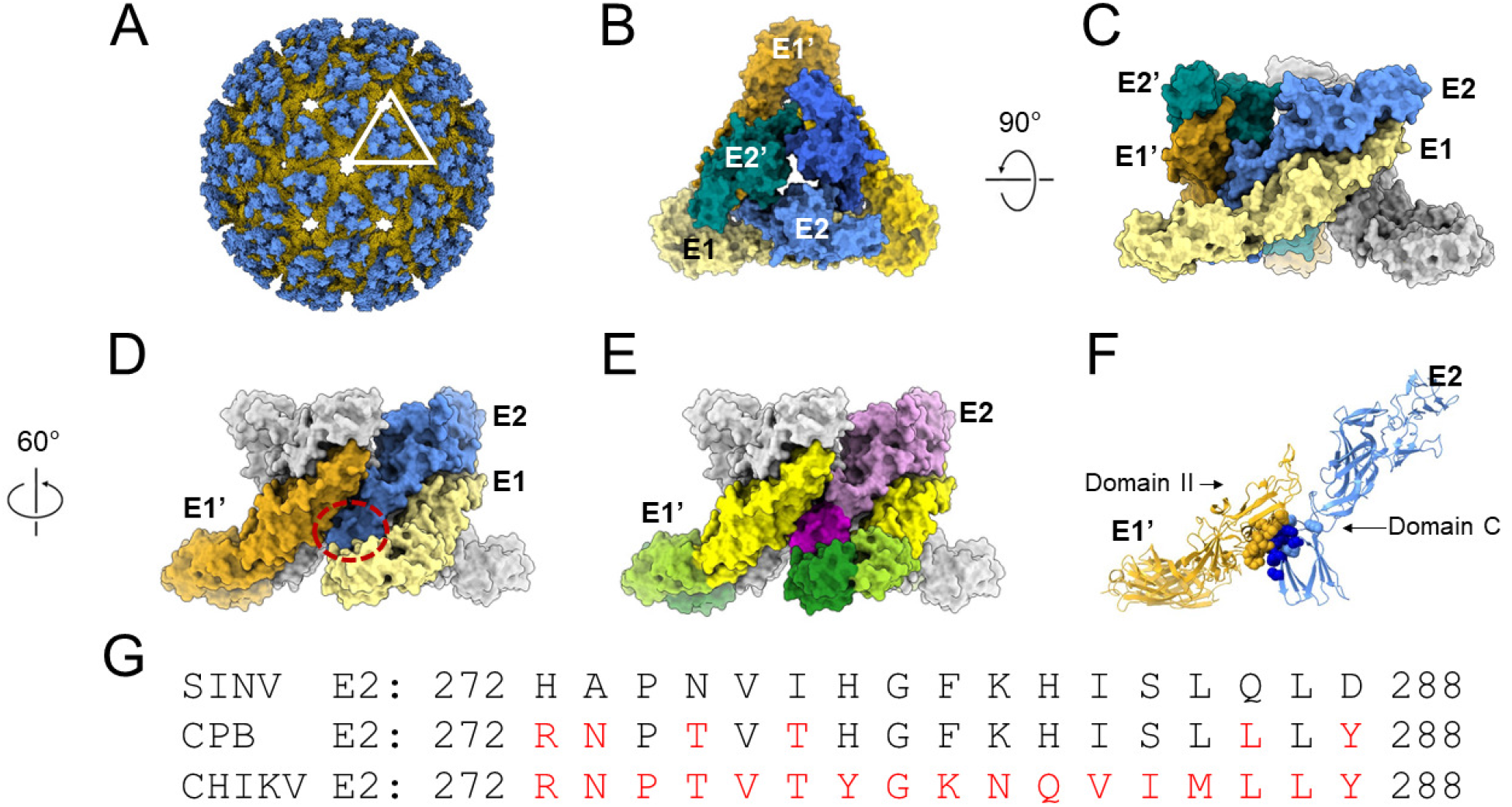
Inter-dimer interface residues in Domain C of E2 may be important for spike assembly. (A) Crystal structure of CHIKV glycoproteins E2 (blue) and E1 (yellow) (PDB ID: 3N40) (29) fit into cryo-EM map of SINV (PDB ID:1Z8Y) (29). One trimeric spike is outlined. (B) During assembly, E1 (*shades of yellow*) forms heterodimers with E2 (*shades of blue*), and these dimers then trimerize. A top view of one of these trimeric spikes, with the individual E2 (in teal, light blue, and blue) and E1 (in gold, light yellow, and saffron) proteins is shown here. A total of 80 spikes are on the surface of the alphavirus particle. (C) The trimer is rotated 90 degrees for a side view. The light blue E2 and light yellow E1 are a heterodimer. Intra-dimer contacts occur between Domains I and II of E1 with Domains A, B, C of E2. The teal E2’ and gold E1’ form another heterodimer, with similar intra-dimer contacts. The last dimer in the spike is shown in grey. (D) A 60-degree rotation of (C) highlights an inter-dimer interface in the trimer. Inter-dimer contacts are between the gold E1’ of one heterodimer and the light blue E2 in the adjacent dimer. Domain C of E2 (red dashed circle) is sandwiched between the adjacent E1’ (gold) and its cognate E1 (light yellow), forming inter-dimer and intra-dimer contacts, respectively. (E) The same proteins, E1’, E2, and E1, shown in (D) are color coded by domain here. Domain III of E1/E1’ is dark green, Domain I of E1/E1’ is yellow-green, and Domain II of E1/E1’ is yellow. Domain C of E2 is in dark purple and Domain I and II are in light purple. The other monomers are colored gray for clarity. (F) Ribbon diagram of the inter-dimer of E1’-E2. Residues in E1’ that contact E2 are in yellow residues in E2 that contact E1’ are in blue spheres (light and dark). The residues mutated in this study are in dark blue spheres. (G) Amino acid alignment of CHIKV and SINV E2 in the inter-dimer region; SINV residues shown in black and CHIKV residues shown in red. The CHIK-Piggy-Back (CPB) chimera was generated by substituting the non-homologous E2 CHIKV residues for the corresponding SINV E2 residues (*highlighted in red in CPB)* in this region.

Voss et al., identified ten residues in E2 Domain C that contact 12 residues in E1 Domain II in the adjacent heterodimer (29). In the low pH SINV E2/E1 heterodimer structure, Li et al identified three residues in E2 Domain C that contact four residues in E1’ Domain II, all also identified in the CHIKV structure (30). Residues E2-272 to E2-288 in Domain C contained a majority of the E2 inter-dimer contacts (Figure 1F, 1G). While there are 13 residues that differ between SINV and CHIK in the primary amino acid sequence in the E2-272 to E2-288 region, only six of these residues are in contact with E1’ in the tertiary structure. Using SINV as our parental virus, we mutated the six residues in this region from SINV to residues found in CHIKV, which belongs to a different clade than SINV (Figure 1F, 1G) (2, 34). The resulting mutant was named CHIK-Piggy-Back (CPB), as the SINV E1’ residues “piggyback” on the six E2 residues that were introduced from CHIKV. We opted to focus on the E2 inter-dimer residues because there are fewer contact residues in E2 than E1 and we speculated that mutating a larger number of residues in E1 would have increased the chance of misfolding due to too many disrupted interactions within the E1 protein itself (21).

### Growth of CPB is attenuated in mammalian cells compared to in mosquito cells

To determine the effect of our CPB mutation, we infected mammalian (BHK-21) and mosquito (C6/36) cells at an MOI of 1 PFU/cell and quantified infectious virus release over time. In BHK cells, the CPB growth was attenuated, producing virus 1 to 1.5 logs lower titer than WT SINV (Figure 2A) as early as 8 hours post-infection. Additionally, CPB had a mean plaque size of 1 mm compared to the mean plaque size of 2 mm for WT SINV (Figure 3A, 3C). In C6/36 cells, however, CPB grew at a similar rate compared to WT SINV (Figure 2B). BHK cells are typically grown at 37°C, while C6/36 cells are grown at 28°C. To rule out temperature dependence on growth, we also measured WT SINV and CPB growth in BHK cells grown at 28°C. We found that there was still a 1 to 1.5 log reduction in CPB titer relative to WT SINV (Figure 2C) suggesting attenuated CPB growth is mainly an effect of different host cell environment, rather than solely different temperatures.

**Figure 2:**
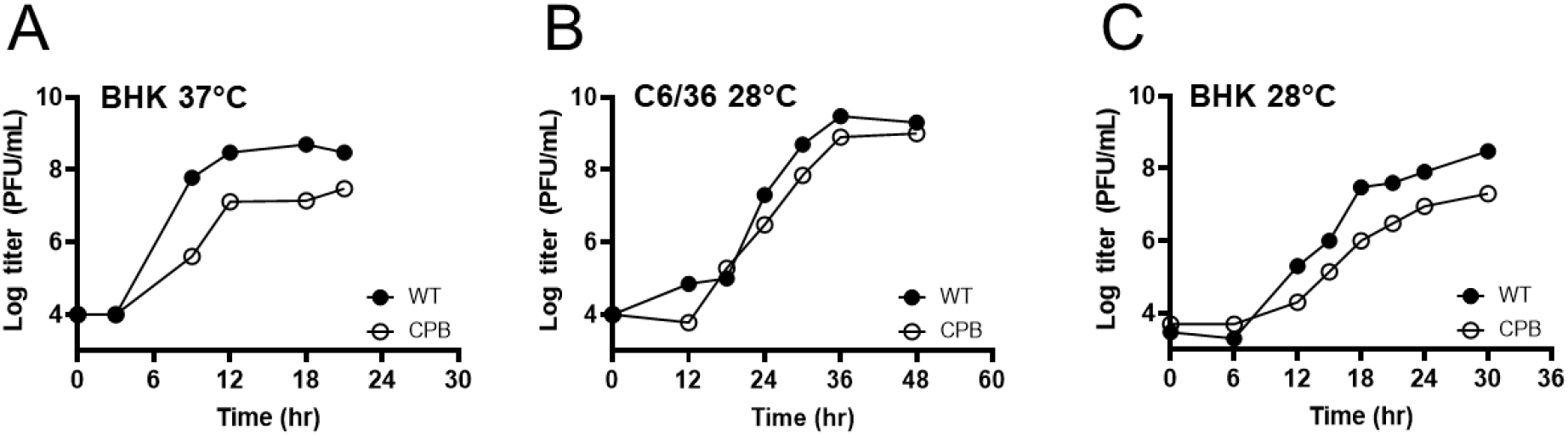
CPB grows slower than WT SINV in mammalian cells compared to in mosquito cells. Cells were infected at an MOI of 1 PFU/cell. At the indicated time points, media was collected and replaced with fresh media. The titers of the collected samples were determined by standard plaque assay on BHK cells. Results are shown for one representative experiment (N=5). (A) Growth kinetics of infectious virus released from BHK cells at 37°C show CPB was attenuated by 1-1.5 logs relative to WT SINV. (B) Growth kinetics of virus grown in C6/36 cells at 28°C show CPB releases infectious particles at the same rate as WT SINV. (C) Growth kinetics of virus grown in BHK cells at 28°C show that CPB growth is still attenuated by 1-1.5 logs relative to WT SINV.

**Figure 3:**
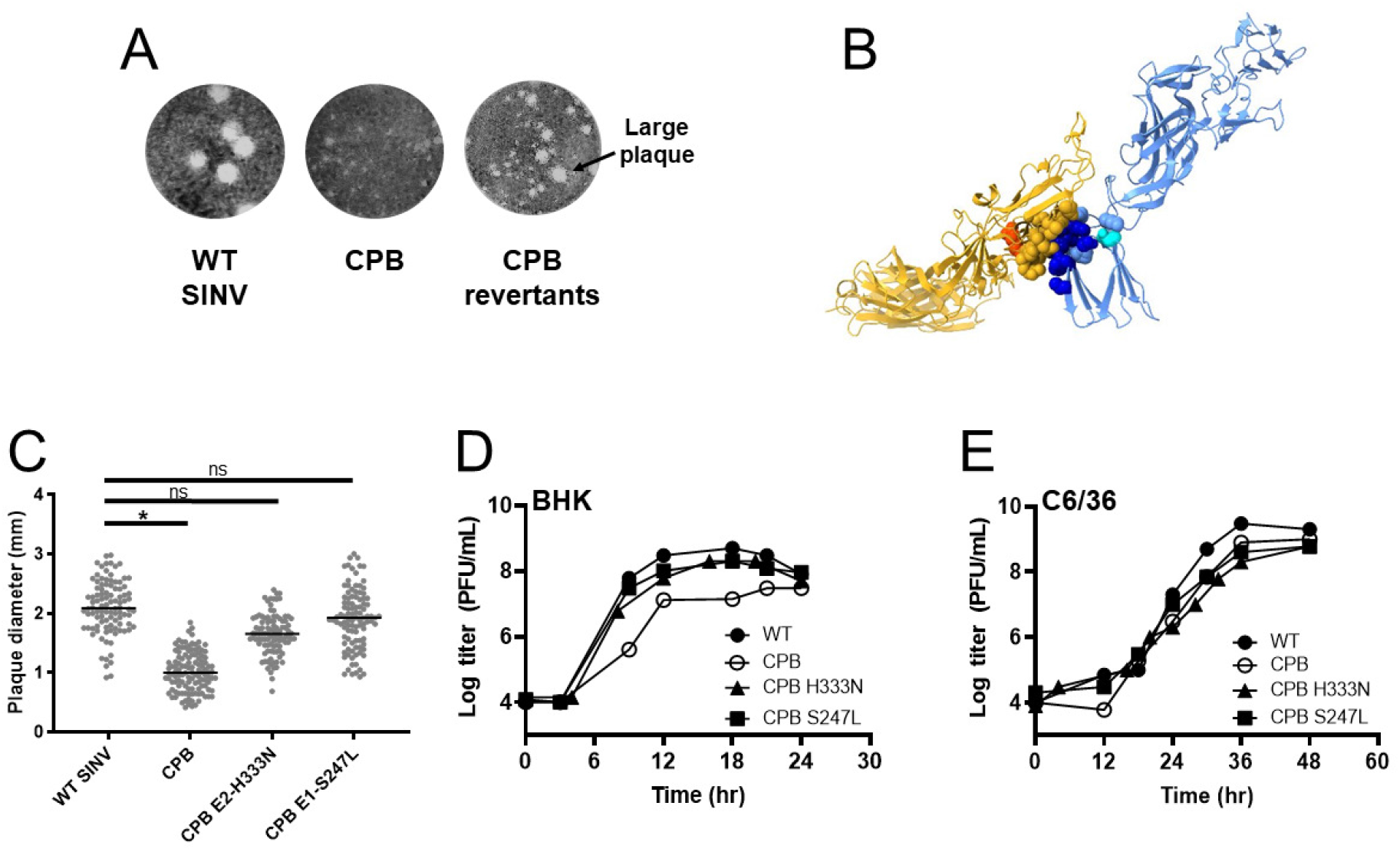
Two CPB revertants map close to the inter-dimer interface and grow similarly to WT SINV. (A) CPB collected 20-40 hours post-electroporation have smaller plaques compared to WT SINV plaques (p<0.0015, see 3C). When CPB is harvested later, or as it is passaged in BHK cells, larger plaques are seen in addition to the small plaques. Five of these larger plaques were isolated, the RNA was sequenced, and two independent second-site revertants were found. (B) The locations of two second-site revertant sites, H333N in E2 (*cyan*) and S247L in E1 (*red)*, are shown in the E1’-E2 dimer. E1’ and E2 contact residues are colored in yellow and blue, respectively, as they were in Figure 1F; mutated sites in CPB are in dark blue. (C) CPB E2-H333N and CPB E1-S247L were introduced independently into CPB. Plaque size of CPB E2-H333N and CPB E1-S247L in BHK cells was larger compared to CPB and not significantly (ns) different from WT. (D) Growth kinetics of the two revertants show they have comparable titers as WT SINV in BHK and (E) C6/36 cells. Representative curves are shown (N=3). Cells were infected at an MOI of 1 PFU/cell, media was collected at the indicated time points, and samples titered on BHK cells.

### Two separate second-site revertants identified for CPB

As we worked with the CPB virus, we noticed that the virus reverted quickly as indicated by the change in plaque size phenotype from small to large. Changes in plaque size were evident often after cells were infected more than 40 hours or passaged more than two or three times (Figure 3A). To isolate revertants, we passaged CPB in BHK cells and saw a mixed plaque phenotype. We isolated and plaque-purified larger plaques, and then isolated and sequenced the viral RNA. We identified two independent single-site revertants, E2-H333N and E1-S247L (Figure 3B). E2-H333N was seen four times, and E1-S247L was seen once. Another mutation, E1-P250S, was seen twice but only in combination with E2-H333N. We focused on E2-H333N and E1-S247L since these single-sites potentially changed the virus fitness on their own. Both the E2-H333N and E1-S247L mutations were in close proximity to the inter-dimer interface (Figure 3B). E2-H333N is spatially near the six-residue cluster we mutated in E2, approximately 13 Å away from the center of the E2 inter-dimer contacts. E1-S247L is approximately 20 Å away from the center of the ten residues of E1’ which interact with the six residues of E2. SINV E1 is glycosylated at position 245. The E1-S247L mutation abrogates the N-X-S/T motif needed for N-linked glycosylation at residue E1 245 (35), by changing the Ser to Leu (Table 1).

**Table 1:** Location of glycosylation sites.

| **Virus** | **Protein** | **Glycosylation site** |
| --- | --- | --- |
| SINV | E2 | N196 |
| SINV | E2 | N318 |
| SINV | E1 | N139 |
| SINV | E1 | N245 |
| CHIKV | E2 | N263 |
| CHIKV | E2 | N345 |
| CHIKV | E1 | N141 |

To determine how these point mutations affect viral assembly, both of these mutations were inserted back into CPB and the WT SINV virus. The plaque sizes of the two revertants cloned into CPB were larger, comparable to that seen with WT SINV virus. The mean plaque size of CPB E2-H333N was 1.6 mm and of CPB E1-S247L was 1.9 mm; both larger than CPB which was 1 mm (Figure 3C). Next, growth kinetics of CPB E2-H333N and CPB E1-S247L were examined independently by infecting BHK cells (Figure 3D). Both revertants resulted in a higher virus yield which grew between 1-2 logs better than the parental CPB and at a similar rate as WT SINV, indicating that both revertants rescued growth in CPB background. We also conducted growth kinetics in C6/36 cells (Figure 3E) and observed no difference between the two revertants in CPB, CPB and WT SINV. When the two mutations were independently cloned into the WT SINV backbone and the viral growth examined in both BHK and C6/36 cells, we observed no change in plaque size or growth kinetics. These results indicate that both of the mutations are neither deleterious nor enhancing in the absence of the CPB mutations.

### CPB particles show defects in spike incorporation compared to WT SINV

Alphavirus spikes initially form dimers and then these dimers trimerize before localizing to the plasma membrane. If the spikes do not trimerize, transport to the plasma membrane is reduced. Spike trimerization is thought to be important for alphavirus budding. In our CPB mutant, we used the alphavirus structures and targeted residues that we thought would disrupt spike trimerization but not dimerization as determined from Voss et al. and Li et al. (29, 30). To determine how particle budding was affected in the CPB mutant, we looked at the composition and morphology of purified virus particles from both BHK and C6/36 cells.

We purified virus particles through a sucrose cushion, ran them on an SDS-PAGE gel, and examined the protein composition. We noticed that BHK purified CPB particles have reduced amounts of E1 and E2 glycoproteins relative to capsid protein when compared to the amounts of these proteins in WT SINV (Figure 4A). This could suggest fewer spikes are associated with the virion and/or the associated spikes may easily dissociate during the purification process compared with WT particles. The E1 protein band in CPB E1-S247L migrates faster than E1 in WT SINV. The E1 protein in CPB E1-S247L is at the same position as the E1 band in the SINV E1-N245/246Q virus which is known to have one of the two E1 glycosylation sites disrupted resulting in a faster migrating E1 protein band. As an additional control, the purified virion of SINV E1-N139Q/E2-N318Q which has one E1 glycosylation site and one E2 glycosylation site disrupted resulting in faster migration in both of those protein bands (Figure 4A) was also included (35).

**Figure 4:**
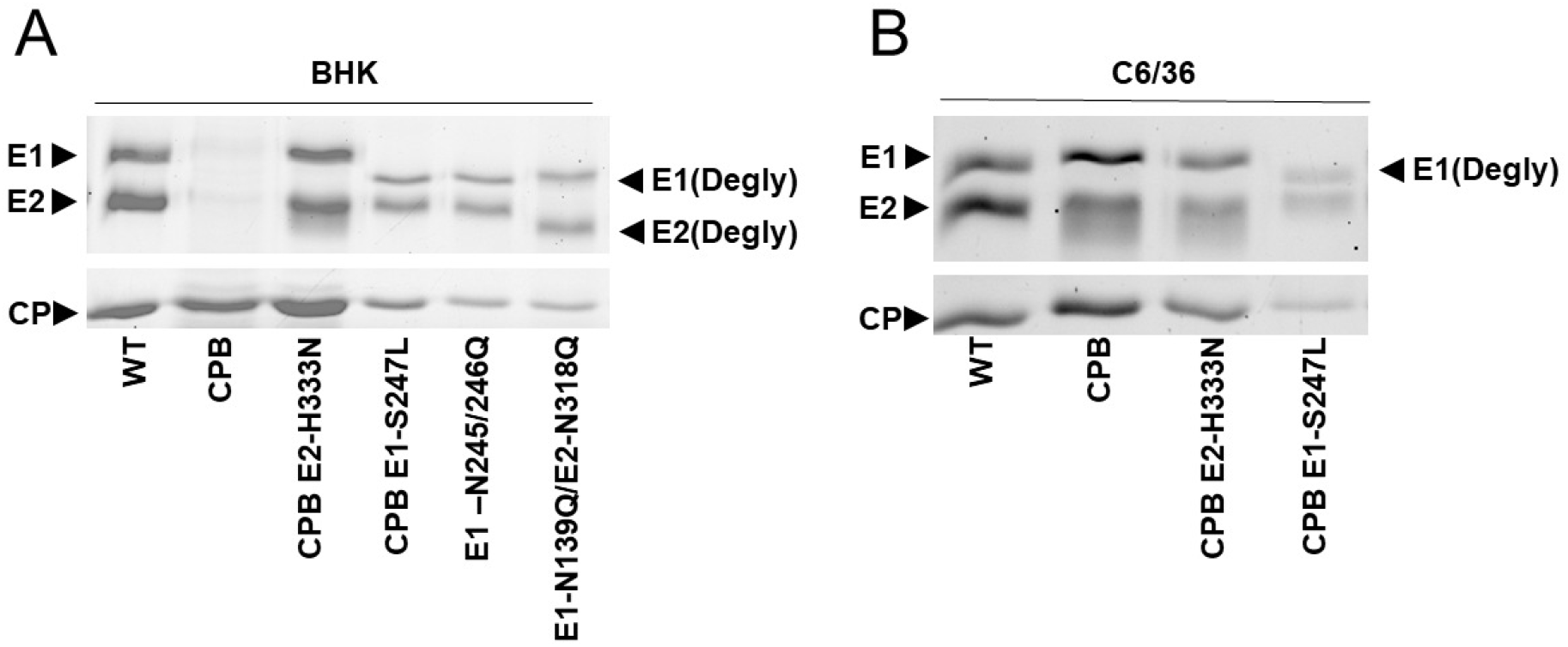
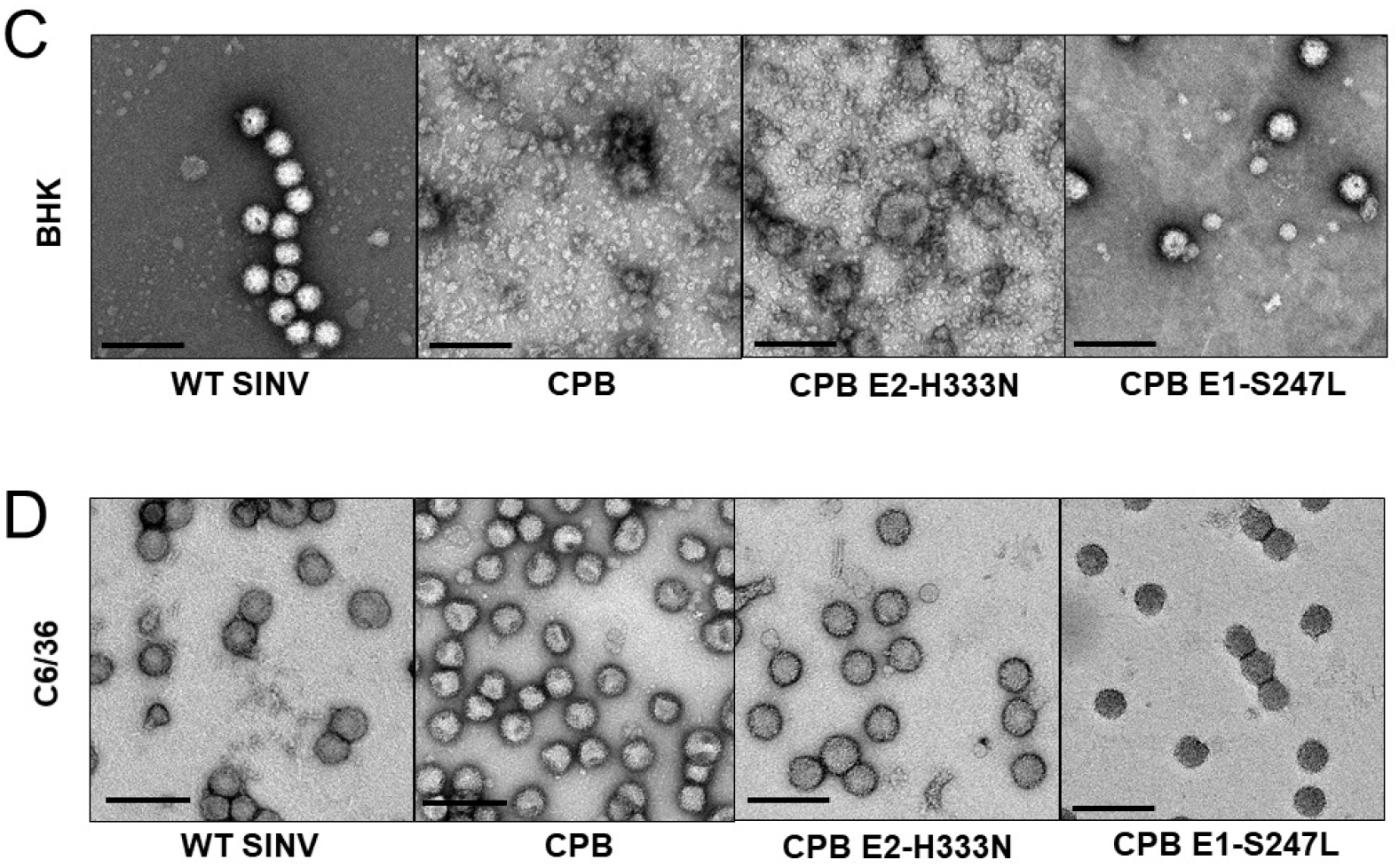
Particle morphology and composition is restored in CPB revertants to varying degrees. Cells were infected at a MOI of 0.1 PFU/cell for 1 hour, after which serum-free media was overlayed. Infected BHK media was collected after 24 hours, and infected C6/36 media was collected after 120 hours. The media was clarified and purified through a sucrose cushion at 140,000 × g for 2.5 h. (A and B) To determine the composition of the virions, purified virus was solubilized in reducing SDS sample buffer, run on an 8% SDS-PAGE gel, and imaged by stain-free TCE. CPB particles have a reduced level of E2/E1 glycoproteins in particles purified from BHK cells. The E1 deglycosylation control is SINV E1-N245/246Q, which has one of its two E1 glycosylation sites deglycosylated resulting in a faster migrating E1 protein band, and the E1/E2 deglycosylation control is E1-N139Q/E2-N318Q and has a deglycosylation site in both E1 and E2 resulting in faster migration in each of those protein bands. (C and D) Purified particles were applied to a Formvar- and carbon-coated 400-mesh copper grid, stained with 2% uranyl acetate, and imaged at ×20,000 magnification with the JEOL 1010 transmission electron microscope. Scale bar for TEM = 200nm.

In contrast to BHK cells, C6/36 purified CPB particles showed similar amounts of E1 and E2 glycoproteins to WT SINV (Figure 4B), suggesting qualitatively that the amount of spike proteins in released and purified particles were similar to WT SINV. The revertants also had similar protein composition to WT SINV with CPB E1-S247L having a faster migrating E1 protein band (Figure 4B). In both CPB and CPB E2-H333N virions, the E2 protein is a smeared band suggesting heterogenous glycosylation on the E2 protein (36).

### CPB particles from mammalian cells show defects in particle morphology, which are partially or fully rescued by revertants

The reduced amount of glycoproteins in CPB particles suggested that the CPB particle may be morphologically different compared to WT particles. To test this, we stained our particles with uranyl acetate and imaged using transmission electron microscopy (TEM). WT SINV is approximately 70 nm in size and spherical and this is observed in both WT particles purified from BHK (Figure 4C) and C6/36 cells (Figure 4D). CPB, however, made almost no identifiable virus particles when purified from BHK cells (Figure 4C). We also noticed that any visible particles were not spherical. CPB E2-H333N partially rescued particle morphology while CPB E1-S247L fully recovered particle morphology (Figure 4C). CPB E2-H333N made more particles than CPB, and some of them looked to be around 70 nm in size; however, many non-spherical particles were still seen. On the other hand, CPB E1-S247L made primarily spherical particles of 70 nm diameter, similar to WT SINV. In both CPB E2-H333N and CPB E1-S247L, there are also small particles ranging in size of approximately 10-40 nm, which could be assembly intermediates or disassembled fragments. These smaller particles suggest that CPB E2-H333N and CPB E1-S247L particles may still be fragile compared to WT SINV, despite being just as infectious.

WT SINV, CPB, and revertant virus particles purified from C6/36 were spherical and homogenous in shape, consistent with no major assembly or growth defects (Figure 4D). CPB particles appeared to have dimples or creases. This could suggest that even in C6/36 cells the mutations made in CPB are detrimental to proper spike formation or have defects in assembly but are still good enough that infectious particle assembly occurs (28).

### Glycoprotein transport is not significantly affected in CPB

The low amounts of glycoproteins in CPB virions compared to WT SINV virions from BHK cells led us to two hypotheses. One, CPB has defects in spike trimerization and transport to the plasma membrane is diminished. Or second, CPB virions are misassembled because inter-dimer contacts have been disrupted. As a result, CPB may disassemble more than WT SINV during the purification process. These options are not mutually exclusive.

We used immunofluorescence to test if glycoprotein transport was altered in CPB compared to WT SINV. We infected BHK cells at an MOI of 2 PFU/cell for ten hours, a time where there was a difference in infectious particles released and probed for glycoproteins at the cell membrane (Figure 5). Cells were fixed with paraformaldehyde, a non-permeabilizing fixative, and probed for SINV glycoproteins. Under these infection conditions, cells were not displaying CPE (Figure 5A, 5C). There was no drastic difference in glycoprotein levels at the plasma membrane between WT SINV, CPB, and revertant infected cells (Figure 5C, 5D). Virus-infected cells were equally healthy and had comparable amounts of detectable glycoproteins.

**Figure 5:**
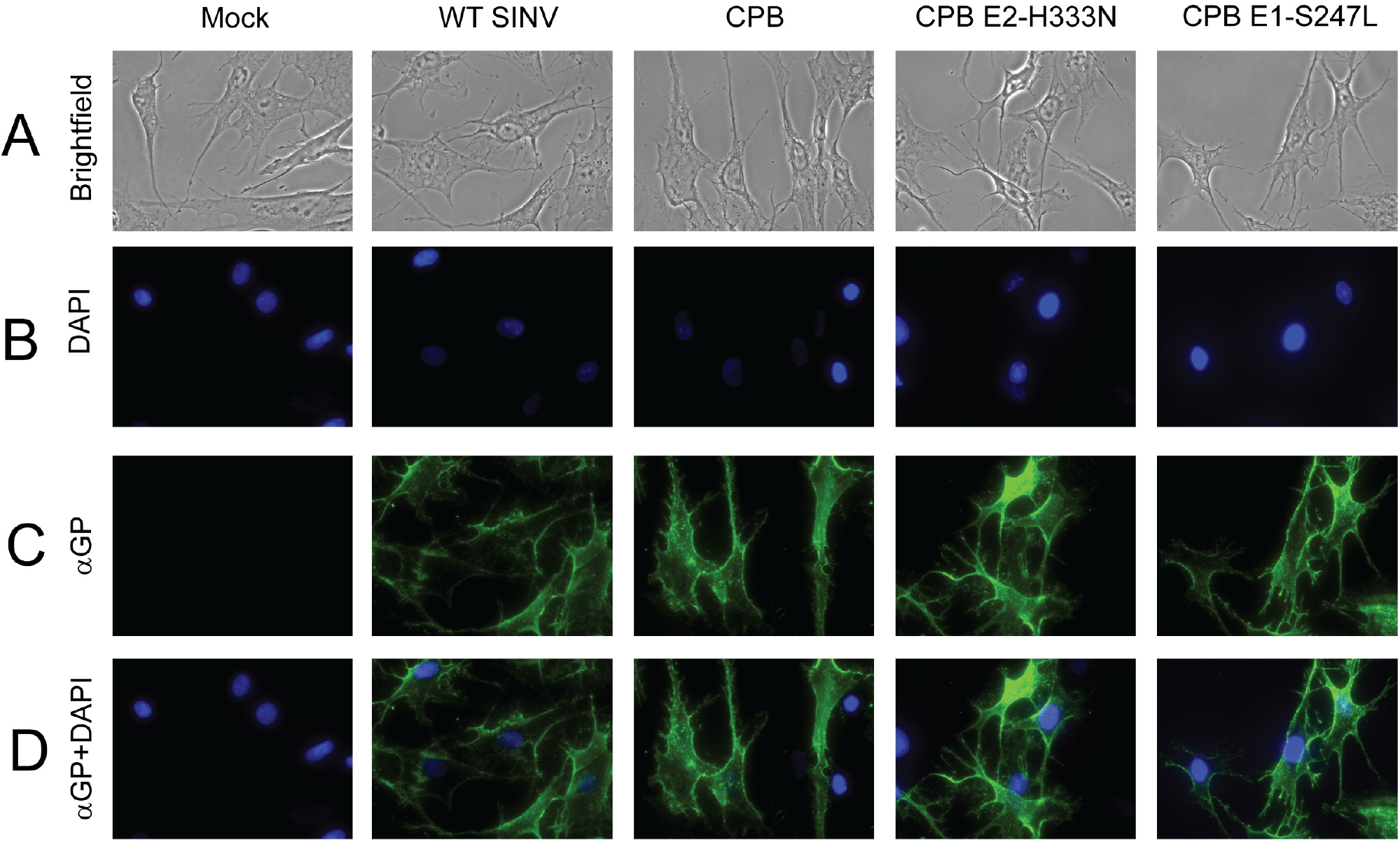
CPB glycoprotein spikes are transported to the plasma membrane of BHK cells in approximately equal amounts compared to WT. BHK cells were infected at an MOI of 2 PFU/cell. Ten hours post-infection, the cells were fixed with 4% EM-grade paraformaldehyde and probed for SINV E1/E2. Images were taken at ×40 magnification on a Nikon Ni-E microscope. Representative images are shown (N=4). (A) Brightfield images and (B) DAPI staining show cells in field of view and their nuclei in blue. (C) Cell surface E2/E1 expression shown in green, Alexa Fluor 488 was the secondary antibody. (D) Anti-E2/E1+DAPI merge shows green fluorescence from surface glycoprotein expression relative to the cells present.

### Spikes in CPB particles are distorted compared to WT SINV

The difference in particle morphology between CPB and WT SINV in mammalian cells was striking when imaged by TEM (Figure 4C, 4D). To further examine the particle structure and spike morphology, we purified virions and used cryo-EM to solve their 3D reconstructions. We focused on WT SINV, CPB, and the revertant CPB E1-S247L since these had the most extreme phenotypes and their structures would be most clear by cryo-EM (Figure 6). We imposed icosahedral averaging (Table 2) during the reconstruction process. The overall structural organizations of all six particles (WT SINV, CPB, CPB E1-S247L from BHK and C6/36) were preserved (Figure 6A-F). There are 80 petal-like spikes arranged into a T=4 surface lattice; the white triangle delineates one asymmetric unit (Figure 6A) (37). No clear E3 density was observed in any of the particles, consistent with what has been previously observed with SINV (8). The glycan modification at E2-196, located at the distal end of the E2 protein, was clearly seen when the estimated resolution is better than 10 Å (red arrows). As seen in other alphavirus structures (7, 38-42), the E1 protein contributes to the continuous shell underneath the spikes (Figure 6A-F, yellow color) with holes at every twofold and fivefold above the membrane. Note the resolution of the particles range from 4.5 Å to 10.7 Å, which reflects both the number of particles used in the reconstruction (Table 2) and the heterogeneity of the particles themselves.

| Reconstruction data | WT SINV<br>(BHK) | CPB<br>(BHK) | CPB E1-S247L<br>(BHK) | WT SINV<br>(C6/36) | CPB<br>(C6/36) | CPB E1-S247L<br>(C6/36) |
| --- | --- | --- | --- | --- | --- | --- |
| **Data collection information** |  |  |  |  |  |  |
| Electron microscope | TFS Titan Krios | TFS Titan Krios | TFS Titan Krios | JEOL 3200FS | TFS Titan Krios | TFS Titan Krios |
| Operation voltage (kV) | 300 | 300 | 300 | 300 | 300 | 300 |
| Electron detector | Gatan K3 | Gatan K3 | Gatan K3 | DE-12 | Gatan K3 | Gatan K3 |
| Energy filter (slit width in eV) | 20 | 20 | 20 | 20 | 20 | 20 |
| Data collection mode | Counting | Super-resolution counting | Counting | Integrated | Counting | Counting |
| Nominal magnification | 105,000x | 64,000x | 64,000x | 80,000x | 64,000x | 64,000x |
| Pixel size in 3D map (Å)<br>(Pixel size in the censor) | 1.68 (0.84) | 2.72 (0.68) | 1.36 (1.36) | 1.9 (1.9) | 2.72 (1.36) | 1.36 (1.36) |
| Total accumulated dose<br>(e<sup>-</sup>/ Å<sup>2</sup>) | 30 | 30 | 30 | 30 | 30 | 30 |
| **Data process statistics** |  |  |  |  |  |  |
| Box size (pixel) | 500 | 300 | 600 | 440 | 300 | 600 |
| Number of particles | 9037 | 15956 | 16133 | 7102 | 63509 | 65053 |
| CTF estimation | CTFFIND4 | CTFFIND4 | CTFFIND4 | CTFFIND4 | CTFFIND4 | CTFFIND4 |
| Data processing software | RELION 3.1 | RELION 3.1 | RELION 3.1 | RELION 3.1 | RELION 3.1 | RELION 3.1 |
| Symmetry imposition | I2 | I2 | I2 | I2 | I2 | I2 |
| B-factor applied (Å<sup>2</sup>) | -545.7 | N/A | -311.4 | -297.3 | N/A | -237.3 |
| Final resolution (Å) | 8 | 10.7 | 5.9 | 10.3 | 10.7 | 4.5 |

**Figure 6.**
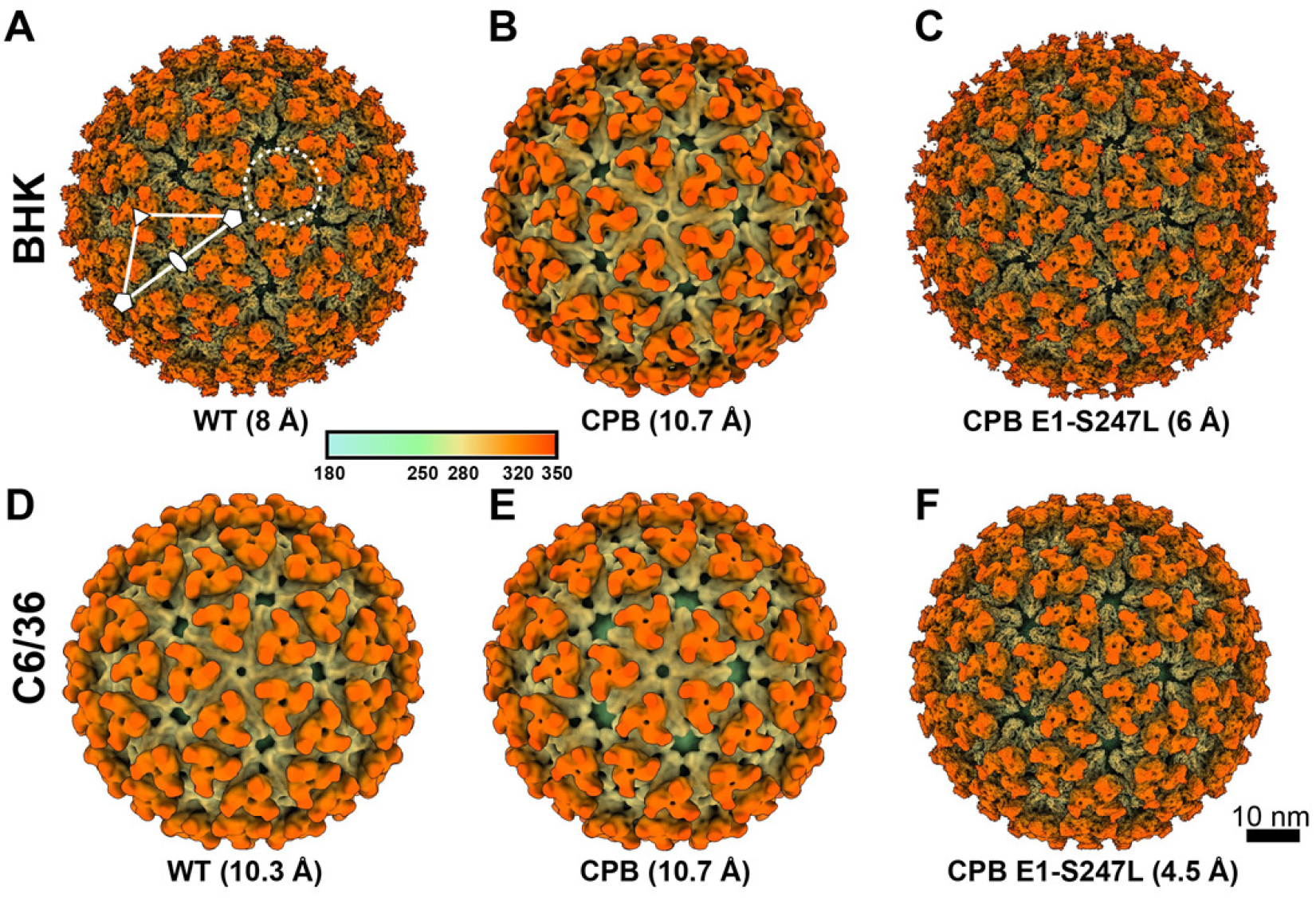
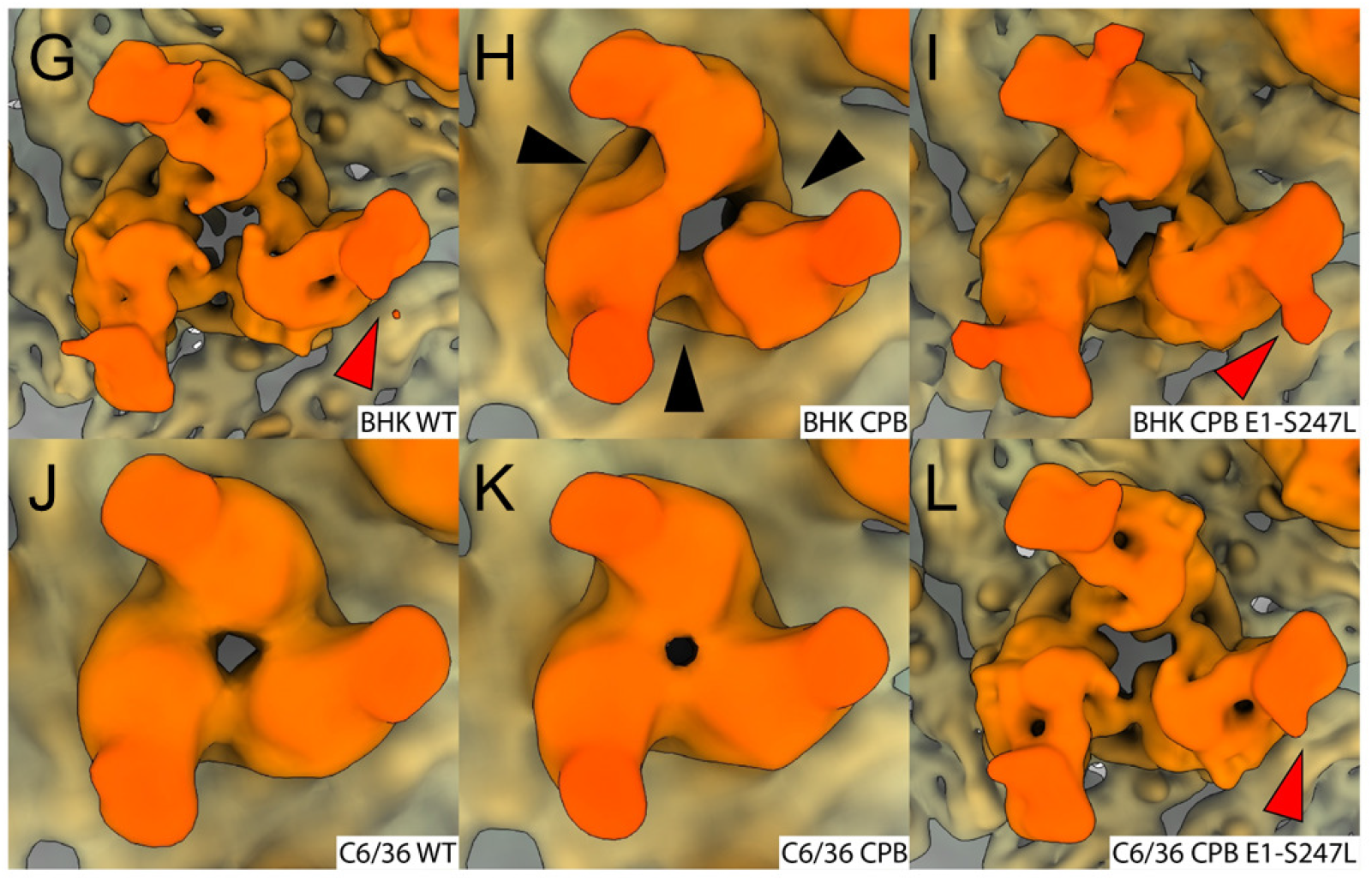
Cryo-EM 3D reconstructions show that CPB spikes from mammalians cells have an altered interdimer organization. (A-F) Radially colored isosurface rendering of WT SINV, CPB, and SINV E1-S247L from BHK cells (A-C, top row) and C6/36 cells (D-F, bottom row). All views are at the 5-fold axis. All six maps have a similar outer appearance with 80 spikes decorating a fenestrated surface. In (A), the white triangle shows one asymmetric unit. Filled oval, triangle, and pentagon indicate locations of twofold, threefold and fivefold axes, respectively. The dotted circle shows one of the spikes on the quasi-three axis. (G-L) Enlarged views of SINV spikes at the quasi-threefold location, all images shown at 10 Å resolution for comparison purposes. The spike of CPB from BHK (H) shows a distinct organization from other SINV spikes. Black arrowheads indicate the difference in the electron density between each interdimer interface. Red arrowheads show glycan modification at E2-196.

However, when looking more carefully at the enlarged view of each spike at the quasi-threefold axis, (dotted circle Figure 6A), there were discernable differences between the samples (Figure 6G-L). We chose the quasi three-fold spike because spikes at the 5-, 3-, 2-fold would, by default of icosahedral symmetry, be identical. The spikes at the quasi symmetry axes would not have any symmetry imposed on them and differences in morphology would be more evident. To best compare the quasi-three spikes, we show them all at 10.7 Å, the resolution of CPB from BHK cells. The irregular density at the inter-dimer location of the spike was clearest in CPB purified from BHK cells (Figure 6H, black arrows). Note that the density of two lobes at the spike petal were fused together, while the other heterodimer within the same spike remained separate (Figure 6H, BHK CPB). For the other five particles, the E1/E2 dimers within a spike were organized in a trimeric manner with true trimeric symmetry.

## DISCUSSION

To investigate the role of the inter-dimeric contacts in assembly, we used a structure-guided approach and mutated six residues in E2 that, in the stable and metastable structures of the spike, are at the interface with the adjacent E1’ protein (Figure 1) (29, 30). We called this mutant CPB. We found that CPB growth is attenuated in mammalian cells compared to mosquito cells (Figure 2). In mammalian cells, CPB glycoprotein expression on the cell surface is similar to WT SINV (Figure 5), but the released particles show CPB particle composition and morphology is defective in mammalian cells while minimally affected in mosquito cells (Figure 4). Our cryo-EM structures of CPB show that CPB particle spikes do not have three symmetrical lobes. Figure 6 shows a visible gap between two of the lobes and the third, which could suggest that two heterodimers are interacting but the third is not. The mutations in E2 disrupt inter-dimer contacts and we hypothesized that CPB growth falls behind WT SINV because of its subsequent assembly defects. However, it is difficult to separate assembly defects from downstream entry impacts.

The difference in phenotype in mammalian and mosquito cells suggests that there is a host factor that is involved in the trimerization of E2-E1 heterodimers since the same viral proteins are assembling in each host cell. If this assembly was autonomous, we would not observe a host-specific difference in both infectious particle production and particle morphology. These differences could be due to different post-translational modifications by the host, different host-protein chaperones affecting virus assembly, or a combination of both. BHK and C6/36 cells are defective some innate immune responses so the attenuation in growth is not entirely a host response to infection. The CPB mutant and its revertants will be useful to identify host chaperones important in spike assembly.

### Stability, composition, and conformation of CPB and revertant spikes

Our immunofluorescence results show that glycoprotein transport to the plasma membrane is comparable between CPB and WT SINV. However, nothing about the conformation of the spikes can be concluded. Furthermore, while E2-E1 are most stable as heterodimers, small amounts of the monomeric proteins will localize to the plasma membrane (15). Our results cannot differentiate conformation or oligomeric state of the glycoproteins at the cell surface.

It is perplexing that we can obtain a structure of CPB purified from BHK cells when this same sample shows few particles by negative-stain TEM and there were low levels of glycoproteins when we look at particle composition on an SDS-PAGE gel (Figure 4). There are several possibilities for these discrepancies, and likely more than one is a contributing factor.

Due to the trimerization defect, fewer spikes may be incorporated into virus particles, and those that are on the particle may be more heterogenous compared to WT SINV. The CPB trimer has a different conformation than the WT SINV trimer, it could also be that the cytoplasmic domain of E2 is no longer able to interact with capsid in a 1:1 ratio and this affects titer and particle stability. The reduced number of spikes on the particles explains the lower titers (Figure 2) and why purified particles show lower amounts of spike proteins in CPB compared to WT SINV (Figure 4). Our cryo EM results also support this hypothesis. We used roughly 16,000 CPB and CPB E1-S247L particles each from BHK infected cells for our reconstructions. Yet the resolutions of the final structures were ∼11Å and 6Å respectively, suggesting more heterogenous particles for CPB than the revertant. WT SINV only used 9,000 particles and a resolution of 8Å was obtained. From our cryo-EM reconstructions, we can see heterogeneity in the individual spikes of CPB compared to WT SINV and CPB E1-S247L particles. Interestingly, our TEM images show that CPB S247L produces particles that are much more WT-like than CPB H333N, despite both revertants restoring CPB growth.

Together the lower number of spikes and their misassembly and heterogeneity could account for particle instability. Fewer spikes on the surface means there is no E1 lattice that covers the viral envelope. The particles are more fragile and sensitive to chemicals. Particles are dehydrated and treated with an acidic stain when preparing for negative stain imaging, and as a result, particles may disassemble or aggregate. In contrast, flash freezing the samples for cryo preservation has a mild effect on the particle’s integrity so misassembled particles will be frozen and can be analyzed.

### Possible implications of revertant residues

CPB quickly reverted resulting in two independently fit mutations, E2-H333N and E1-S247L, and one mutation, P250S, which was seen only in combination with E2 H333N. E2-H333N, E1-S247L, and P250S are all in close proximity to the cluster of mutations made at the inter-dimer interface, although none were identified by Voss or Li as one of the inter- or intra-dimer contact residues (29, 30). Residue 333 for most viruses is either a histidine or an asparagine. Additionally, the corresponding residue for SINV H333 is N330 in CHIKV. Because the histidine side chain has a pKa of 6.00, it is within the range of different pH environments in the host cell. This histidine residue may affect CPB either during entry, specifically during disassembly in the endosome (43, 44), or during assembly as the E2-E1 heterodimer travels through the ER and Golgi (45). The role in disassembly could be unfavorable in CPB, since it may have a less stable trimer and less stable heterodimers even without a low pH environment. During assembly of the CPB E2-E1 heterodimers, there may be improper folding possibly making trimer assembly more sensitive to pH which cannot occur with H333.

S247L interferes with N-linked glycosylation motif that allows E1-N245 to be glycosylated. In CHIKV, there is only one glycosylation site on E1 at N141 versus two in SINV, one at N139 and one at N245 (Table 1) (35, 38). Although residue 247 is not conserved among alphaviruses, it is clear that some viruses including SINV have the N-X-S/T glycosylation motif at N245, while others including CHIKV do not. It is interesting that CPB S247L appears more WT-like in particle morphology than CPB H333N. It is possible that de-glycosylation at this site allows for better folding or removes a steric hinderance that allows for trimerization.

From our revertant sequencing we also saw that in some instances when E2-H333N was mutated, E1-P250S was also mutated. Residue 250 is a proline residue in SINV and a serine residue in CHIKV. It was seen with H333N but does not seem necessary for restoring growth since H333N was seen alone two times and CPB H333N independently restored growth. However, the proline residue in CPB might disrupt folding since it is unable to form hydrogen bonds. This could explain why CPB H333N has less spherical looking particles, despite having growth comparable to that of WT SINV. In all three cases, it appears that CPB is reverting to be more like CHIKV.

### Other revertants in spike proteins that show host specificity

Our work is not the first time a mutation has shown host specificity in alphaviruses. In CHIKV, Ashbrook et al., showed a mutation at E2 G82R enhanced infectivity in mammalian cells but reduced infectivity in mosquito cells and reduced virulence in mouse model (46). This residue is present on the exterior of the E2 protein and was determined to be important to GAG binding, entry and virulence.

Jupille et al. identified Ross River E2 Y18H as having a fitness advantage in mosquito cells and a disadvantage in mammalian cells (47). This residue lies in the intra-dimer interface of the E2-E1 heterodimer. Interestingly, in the Ross River clade, the viruses have either a tyrosine or a histidine at position 18 and the authors suggest, this residue serves as a regulator of fitness between the mosquito vector and mammalian host (47). In our work, the revertant E2 H333N was isolated. Most alphaviruses are either a histidine or asparagine at this residue emphasizing the structural and functional requirement of this residue.

Our work presented here is the first example of a mutation that has host specificity and mapping to defects in spike assembly. Further work needs to be done to dissect the exact mechanism of where in the spike assembly pathway the CPB mutant fails in mammalian cells and how the revertants overcome these defects. Presented results now allow us to further identify host-specific chaperones and factors necessary for alphavirus glycoprotein folding and oligomerization and possibly extend to other arboviruses that assemble in multiple host environments.

## Materials and Methods

### Viruses and cells

The virus strains used in this work were the TE12 strain of SINV and 181/21 strain of CHIKV (a gift from Dr. Terrence Dermody). BHK-21 cells (BHK) (American Type Culture Collection, Manassas, VA) were grown in minimal essential medium (Mediatech, Manassas, VA) supplemented with 10% fetal bovine serum (FBS; Atlanta Biologicals, Lawrenceville, GA), nonessential amino acids, L-glutamine, and antibiotic-antimycotic solution (Corning, Corning, NY). BHK-21 cells were grown at 37°C in the presence of 5% CO_2_, or at 28°C in the presence of 5% CO_2_. C6/36 cells (American Type Culture Collection, Manassas, VA) were grown in identical medium at 28°C in the presence of 5% CO_2_.

### Generation of wild-type and mutant viruses

The CPB mutant virus was generated using a two part QuikChange site-directed mutagenesis (Agilent, Santa Clara, CA) of the TE12 SINV cDNA clone. First the residues between nucleotides 9444 and 9494 were deleted, and then the chimera sequence was inserted. No additional changes were introduced to the chimera virus during cloning. The mutations were confirmed by sequencing the E2 region. The CPB E2-H333N and CPB E1-S247L were generated by QuikChange site-directed mutagenesis.

Wild-type and mutant cDNA clones were linearized with Sac I and in vitro transcribed with SP6 polymerase at 39°C for 2 hr. For electroporation of BHK cells, approximately 10^7^ BHK cells were trypsinized, washed two times with phosphate-buffered saline (PBS), and resuspended with PBS to a final volume of 500 μl. The cells were mixed with *in vitro*-transcribed RNA in a 2-mm-gap cuvette and pulsed once at 1.5 kV, 25 μF, and 200 Ω using a Bio-Rad Gene Pulser Xcell electroporation system (Bio-Rad Laboratories, Hercules, CA). Following a 5-min recovery at room temperature, the cells were diluted 1:10 in cell medium and incubated at 37°C in the presence of 5% CO_2_. At the indicated time points (around 24 hours post infection for WT, CPB E2-H333N, CPB E1-S247L, and 40 hours post infection for CPB), virus was harvested, and the titer was determined using a standard plaque assay procedure (23). Plaques were detected at 48 hours post-infection by formaldehyde fixation and crystal violet staining.

### One-step growth analysis

Confluent 12-well plates of BHK cells were infected with virus at MOI of 1 PFU/cell at room temperature for 1 hr. Following this adsorption period, the cells were washed with PBS to remove any unbound particles, and 400 μL of media was added. At the indicated time points, 400 μL media was removed and replaced with fresh media. The titers of these samples were determined by standard plaque assay and plotted against time. *P* values were calculated using Welch’s unpaired *t* test.

### Identification of second-site revertants

Large plaques were detected from media of CPB infected BHK cells more than 40 hours post-electroporation. As previously described (23), large plaques were isolated and used to infect BHK cells. The infected cells were lysed at 14 hours post-infection with TRIzol reagent (Invitrogen, Carlsbad, CA), and cytoplasmic RNA was isolated using chloroform extraction. The coding sequence of the structural polyprotein was amplified from the viral RNA using RT-PCR and two primers, one specific for the E1 region and the other specific for the E2 region. The region corresponding to the structural polyprotein was sequenced to identify the location of the potential second-site mutation. To verify that the large plaque phenotypes were due to the mutation identified in sequencing, the revertant sites were introduced back into the chimeric viruses and growth kinetics and plaque sizes were analyzed.

### Virus purification

150 mm dishes were infected with 3 mL virus at a MOI of 0.1 PFU/cell at room temperature for 1 hr. After the adsorption period, the cells were overlaid with 15 mL serum-free media (Thermo Fisher Scientific life technologies, Waltham, MA) supplemented with nonessential amino acids, L-glutamine, and antibiotic-antimycotic solution (Corning). Medium was collected approximately 24 hours post infection for BHK cells. For C6/36 cells, 5 mL fresh serum-free media was added after 72 hours, and medium was collected after approximately 120 hours post infection. The medium was then spun down at 1157 × g for 5 min at 15°C to remove cells and cell debris. Virus particles from the clarified medium was pelleted through sucrose cushion. The clarified medium was overlaid onto 3 ml 27% sucrose in 20 mM HN buffer (20 mM HEPES, pH 7.5, and 150 mM NaCl) and spun at 140,000 × g for 2.5 h at 15°C (36, 48).

### SDS PAGE gel of purified virus particles

Purified virus particles were loaded onto an 8% SDS gel containing 0.5% 2,2,2-trichloroethylene v/v in the resolving gel, and run for approximately 45 minutes at 200 V. The gel was imaged with the Bio-Rad ChemiDoc MP Imaging System using the stain-free image setting. The PageRuler prestained protein ladder (Thermo Fisher Scientific-Invitrogen, Waltham, MA) was used in all studies.

### Immunofluorescence analysis of cell surface spike protein expression

SINV was purified from BHK and C6/36 cells. The purified particles were ran on an SDS-PAGE gel and the glycoprotein bands were excised and sent to Cocalico Biologics, Inc (Stevens, PA) to generate polyclonal antibodies. The primary antibodies were pre-cleared to reduce background signal from non-specific cell binding. A confluent well of BHK cells was washed twice with PBS, chilled with PBS for 2 hours, and then incubated at 4°C with 200 μL of a 1:10 dilution of antibody in cold PBS for 2 hours while rocking. The antibody mixture was removed and centrifuged, and the supernatant was saved and used in immunofluorescence studies.

BHK cells were grown on coverslips and at approximately 75% confluency were infected with virus at an MOI of 2 PFU/cell at room temperature for 90 minutes. After this adsorption period, fresh media was added, and cells were incubated at 37°C in the presence of 5% CO_2_. Ten hours post infection, the cells were washed with PBS and then fixed with 4% paraformaldehyde (Thermo Fisher Scientific Life Technologies, Waltham, MA) at room temperature for 10 minutes. 16% EM-grade paraformaldehyde was diluted in PBS to make 4% paraformaldehyde; the EM-grade paraformaldehyde is methanol-free to avoid permeabilization. The cells were then washed, blocked in 2.5% BSA in PBS for 30 minutes, and incubated with pre-cleared polyclonal anti-E2/E1 (1:50 of pre-cleared) in 2.5% BSA for 45 minutes. Cells were washed and incubated with Alexa 488 Goat anti-Rabbit secondary antibody (1:5000) in 2.5% BSA in PBS for 45 minutes in the dark. Cells were then washed and stained with DAPI. The coverslips were carefully removed, dipped in distilled water and blotted, and inverted onto 5 µL of Aqua-Poly/Mount (Polysciences, Warrington, PA) on a slide. Slides were imaged using an Olympus 1×71 fluorescence microscope (Olympus, Center Valley, PA).

### Transmission electron microscopy

Four μL of purified virus was applied to a Formvar- and carbon-coated 400-mesh copper grid (Electron Microscopy Sciences, Hatfield, PA) for 25 seconds, washed with 4 μL water for 25 seconds, and stained with 2% uranyl acetate for 25 seconds. The stained grids were analyzed using a JEOL 1010 transmission electron microscope (Tokyo, Japan) operating at 80 kV. Images were recorded using a Gatan UltraScan 4000 charge-coupled-device camera (Pleasanton, CA).

### Cryo-EM imaging and 3D reconstruction

To prepare a frozen-hydrated cryo-EM specimen, approximately 4 μL of purified virus sample was applied to a glow-discharged 300-mesh copper grid coated with continuous carbon film (Electron Microscopy Sciences, Hatfield, PA). The grid was plunged into a liquid ethane container that is further cooled by a liquid nitrogen bath. This process was performed using Vitrobot Mark IV under 4 degree and 100% humidity (Thermo Fisher Scientific-Invitrogen, Waltham, MA). Frozen-hydrated cryo-EM grids for all samples except SINV WT (C6/36) were clipped into cartridges and then transferred into a cassette according to the manufacture protocol. The cassette was loading inside a 300-kV Titan Krios G3i equipped with Gatan BioContinuum™ K3 direct electron detection camera. Data acquisition was set up using TFS EPU under counted super-resolution mode. The nominal magnification is 64,000x (equal to 0.7 Å per pixel) and the illumination has a dose rate of 1 e^-^/Å^2^ per frame under the exposure preset with a total dose of 30 e^-^/Å^2^. Zero-loss peak was aligned every hour with an energy slit opened at 20 eV. Data collection for SINV WT (C6/36) was done by using a 300-kV JEOL JEM-3200FS TEM equipped with DE-12 CMOS camera (Direct Electron). The frozen hydrated grid was transferred to a Gatan 626 cryo-holder and inserted into the TEM. The nominal magnification was set to 80,000x (equal to 1.9 e^-^/Å^2^). The zero-loss peak was aligned at the beginning of the data collection with a slit opened at 20 eV. The total accumulated dose is ∼30 e^-^/Å^2^.

Image analysis was performed using Relion (v3.1) (49). Initial particle picking was done by Laplacian-of-Gaussian filtered auto-picking method implemented in Relion (50). Subsequently, 2D classification was used to eliminate the noise density that got picked earlier and the 3D initial model was built de novo. This initial model was then used as a template for automatic particle picking for all samples. Similarly, multiple runs of 2D classification and 3D refinement were performed to obtain the final 3D models (summarized in Table 2). Each volume was rendered using UCSF ChimeraX (51).

## Acknowledgements

We thank IU Virology group for constructive discussions and Jim Powers, Walczak and Shaw labs and the IU Light Microscopy Facility. Funding for this work from IU Wells Scholars Program and LS McClung Fellowship (SCR), the startup fund from The Pennsylvania State University College of Medicine to J.C.-Y.W and IU IAS (SM). We also gratefully acknowledge The Pennsylvania State University College of Medicine for access to the Cryo-EM (RRID:SCR_021178) and the HPC core facilities.

## Notes

### Competing Interest Statement

The authors have declared no competing interest.

